# Sequencing SARS-CoV-2 in a malaria research laboratory in Mali, West Africa: the road to sequencing the first SARS-CoV-2 genome in Mali

**DOI:** 10.1101/2021.05.05.442742

**Authors:** Antoine Dara, Bouréma Kouriba, Amadou Daou, Abdoul Karim Sangare, Djibril Kassogue, Charles Dara, Abdoulaye Djimde

## Abstract

Next generation sequencing (NGS) has become a necessary tool for genomic epidemiology. Even though the utility of genomics in human health has been proved, the genomic surveillance has never been so important until the COVID 19 pandemic. This has been evidenced with the detection of new variants of SARS-CoV-2 in the United Kingdom, South Africa and Brazil recently using genomic surveillance. Until recently, Malian scientists did not have access to any local NGS platform and samples had to be shipped abroad for sequencing. Here, we report on how we adapted a laboratory setup for *Plasmodium* research to generate the first complete SARS-CoV-2 genome locally. Total RNA underwent a library preparation using an Illumina TruSeq stranded RNA kit. A metagenomics sequencing was performed on an Illumina MiSeq platform following by bioinformatic analyses on a local server in Mali. We recovered a full genome of SARS-CoV-2 of 29 kb with an average depth coverage of 200x. We have demonstrated our capability of generating a high quality genome with limited resources and highlight the need to develop genomics capacity locally to solve health problems. We discuss challenges related to access to reagents during a pandemic period and propose some home-made solutions.

## Background

SARS-CoV-2 virus was first detected in Wuhan, China in December 2019. The SARS-CoV-2 disease also known as COVID-19 has become pandemic and continues to be a global burden impacting all level of health care systems and economies. As of May 3, 2021, over 153 million people have been infected worldwide with more than 3.2 million deaths (https://covid19.who.int/). In sub-Saharan Africa, COVID-19 cases are relatively low compared to other parts of the world even though the reported numbers might be underestimated.

In Mali, the first case of COVID-19 was detected on March 26, 2020. By May 3, 2021, 13,937 cases were reported with 491 deaths according the WHO report. The Malian government implemented protection and control measures of the disease. The country has currently experienced its third wave of the outbreak.

Global efforts to control the disease included genomic surveillance as a key component of control strategies. Genomic epidemiology of COVID 19 has provided a framework for tracing the origins of the circulating virus, identifying clusters or hotspots of transmission, route of transmission, and detecting new variants [1]–[3]. For instance, new variants reported in the United Kingdom, South Africa and Brazil have been identified through genomic surveillance [4] [5], [6], [7].

However, Africa has been lagging behind with providing sequences of the virus. The first African SARS-CoV-2 was provided by Nigeria [8]. So far, few laboratories in Africa are able to sequence the genome of the virus. Most of the samples are usually sent abroad in institutions with sequencing capacities. For instance, the first sequences from Mali deposited in GISAID were sequenced in Germany. However, shipping samples overseas is not only expensive but ethical and regulatory issues constitute major challenges. Therefore, developing local capacity in sequencing seems an optimal option for countries like Mali.

As the pandemic progresses, more sequences are needed to monitor the evolution of the virus. Until recently, Malian researchers did not have access to next generation sequencing locally. The Malaria Research and Training Center at the, University of Science, Techniques and Technology of Bamako has recently acquired a MiSeq instrument for its malaria genomics research. However, the urgency of the COVID-19 pandemic prompted the use of the existing resource to sequence SARS-CoV-2 genome. Indeed, a combination of factors including lack of preparedness, lack of specific funding, difficulty in shipping samples abroad led to the necessity to use malaria research resources to sequence SARS-CoV-2 genomes. We describe here the first genome of SARS-CoV-2 sequenced locally and highlight the importance of implementing genomic surveillance and discuss some challenges related to access to reagents during a pandemic.

## Methods and materials

### Samples

The sample was collected in January 2020 by the Centre d’Infectiologie Charles Merieux’s mobile laboratory which was dispatched by the Ministry of Health to handle an outbreak of SARS-Cov-2 in Timbuktu, more than 1000Km North of Bamako, the Capital city of Mali. Nasal/pharyngeal specimen served as starting material for RNA extraction. Total RNA was extracted Qiamp viral rna mini kit (Qiagen Cat. 57704) according to manufacturers’ instructions. Real-time transcriptase polymerase chain reaction (RT-PCR) was used for the detection of positive samples. RT-PCR tests were based on the N gene and the RNA-dependent RNA polymerase (RdRp) gene located on the ORF1ab.

### Library preparation

RNA was quantified with a fluorometer (Denovix QFX) using Denovix RNA quantification kit. Total RNA was depleted of ribosomal RNA prior to library preparation. Briefly, 200ng of input RNA was subjected rRNA depletion according to the Ribo-Zero Gold rRNA depletion protocol (Illumina, 96 samples, Cat no. 20020599). Libraries were prepared from the depleted RNA using the Illumina TruSeq Stranded Total RNA Library Prep Kit (Cat no. 20020599). The protocol included the following steps: RNA fragmentation, cDNA synthesis, adenylation of the 3’-end, adapter ligation, and cDNA enrichment (TruSeq Stranded Total RNA protocol). For the cDNA synthesis, because no one on campus had the transcriptase Superscript II we slightly modified the Illumina protocol by replacing it with the SuperScript IV (introgen cat. 12594025) and the PCR reaction was performed at 50°C instead of 42°C. cDNA was quantified with the fluorometer using DeNovix dsDNA Broad Range Quantitation Reagent according to the manufacturer’s instructions.

### Library size estimation of the cDNA library using an agarose gel

During this pandemic, access to reagents was a real challenge. While, our order for Bioanalyzer had been pending for months, we decided to estimate our library size using 1% agarose gel. Briefly, 2 ul of the library was loaded using NEB loading dye (Cat: NEB #B7025) because it did not make a shadow on the gel as compared to the bromophenol blue dye. Gel image was visualized and saved as a .jpeg file. The image was subsequently analyzed using a gel analysis tool ImageJ software [9].

### Sequencing

Subsequently, libraries were quantified and normalized with nuclease-free ultrapure water, and spiked with 5% of Phi X control, and run onto the Illumina MiSeq instrument (Illumina^®^, San Diego, CA) for sequencing. MiSeq reagent kit v3 (150-cycle) was used according to the manufacturer’s instructions. Twenty (20) pM of the normalized library was loaded onto the MiSeq instrument.

### Data analysis

All data was processed locally using a recently built bioinformatics infrastructure located at the Malaria Research and Training Center. Raw data (FastQ files) were transferred from the MiSeq instrument (via BaseSpace account) to the local server and analyzed according to the workflow in supplemental figure 1 (Figure S1). Quality control was performed using *FastQC* and *Multi-FastQC* tools. The QC plot showed that all reads were of high quality and did not require any trimming. Subsequently, reads were mapped onto the SARS-CoV-2 reference genome (accession number: MN908947.3) using the Burrows Wheeler aligner (BWA) version 0.7.17 [10] with default parameters. Post-mapping file generated as bam file was processed with *samtools* [11] and the *vcfR* package to determine the percent of aligned reads, the depth of coverage and mapping quality. Library estimation was performed using the CollectInsertSizeMetrics tool version 2.8.1 implemented in Picard. The alignment was visualized with IGV (Interactive Genome Viewer) tool [12]. A consensus sequence algorithm implemented in IGV was also used to generate the consensus sequence from the alignment [13]. *Vcftools* was used to generate a variant call file (vcf). *SnpEff* implemented on Galaxy server as well as Nextclade were used to annotate variants.

**Figure 1:**
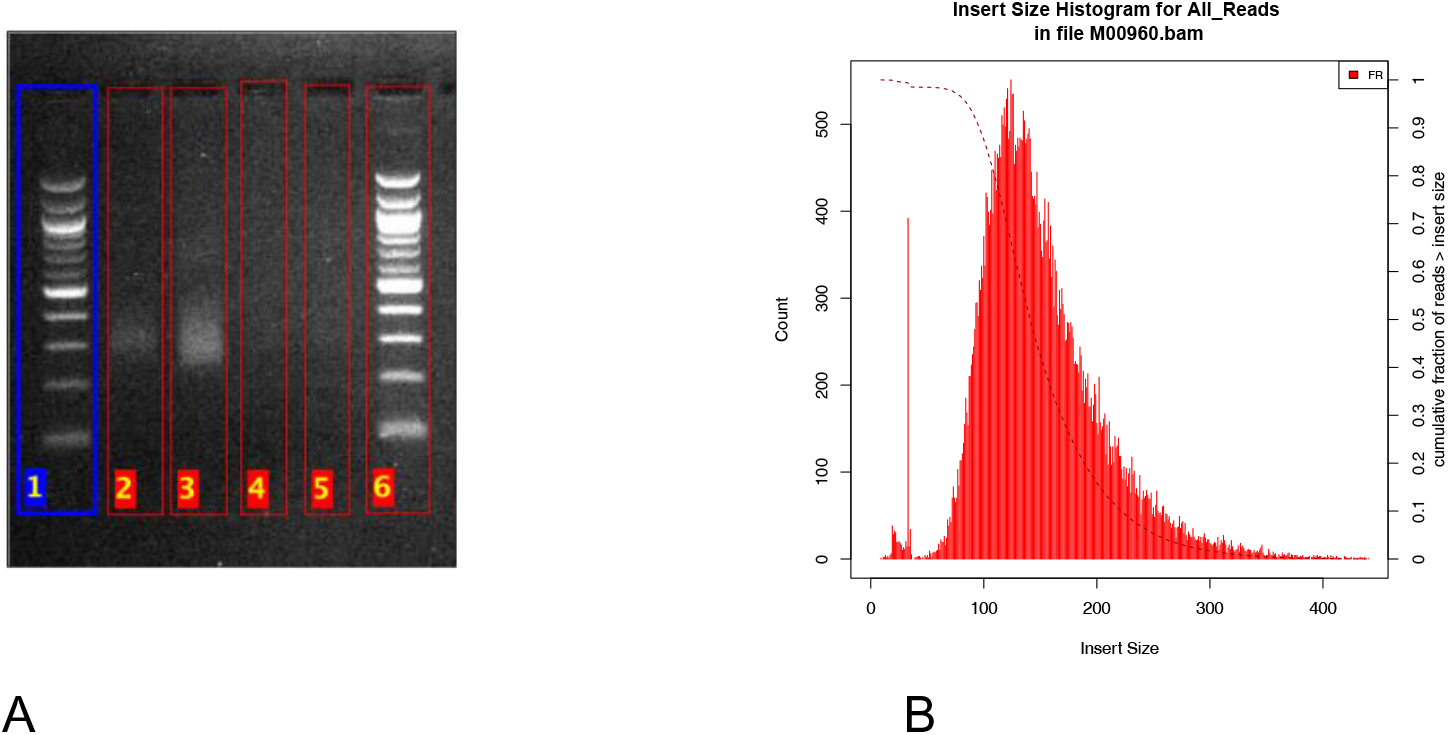
Library insert size estimated from the reads with Picard tool and agarose gel image analysis.

We performed *de novo assembly* of the reads using MEGAHIT software [14]. Shorter contigs (<2000bp) were filtered out and the remaining contigs were queried with BLASTN on the NCBI database to recover the SARS-CoV-2 genome. Consensus sequence from the alignment approach was compared to the contigs from de novo assembly using *nucmer* aligner [15].

## Results

The library size was estimated to around 325 bp on the agarose gel (Figure 1A). After sequencing, the average insert size was estimated from the reads to around 150 bp (Figure 1B).

A total of 78688878 reads (forward and reverse) were obtained from the run. The MiSeq run output generated 6.45 Gbp as raw data with 96.22% of reads having a quality score Q>30 (less than 1 error in every 1000 bp). The average quality measured as Phred-score of the raw data was ~35. Pre-processing of the raw data showed all reads were of high quality (Figure S2). The reads did not require any end trimming.

After pre-processing the data, to confirm that the reads contain SARS-CoV-2 sequences, we mapped the raw reads to the reference Wuhan genome (accession number: NC_045512). Post-alignment processing of the BAM file resulted in 0.13% of the total reads (104479 reads) aligned to the reference genome covering 99% of genome. An average read depth of 250 (Figure 2) and a mapping quality > 50 were obtained (Figure S3). Both the forward and the reverse aligned well to the reference genome (Figure S4). We generated a consensus genome of approximately 29 903 base pairs from the bam file using a consensus mode implemented in the IGV tool. BLAST analysis showed the top hit with 99.95 percent identity covering 100% of the query sequence (Figure S5).

**Figure 2:**
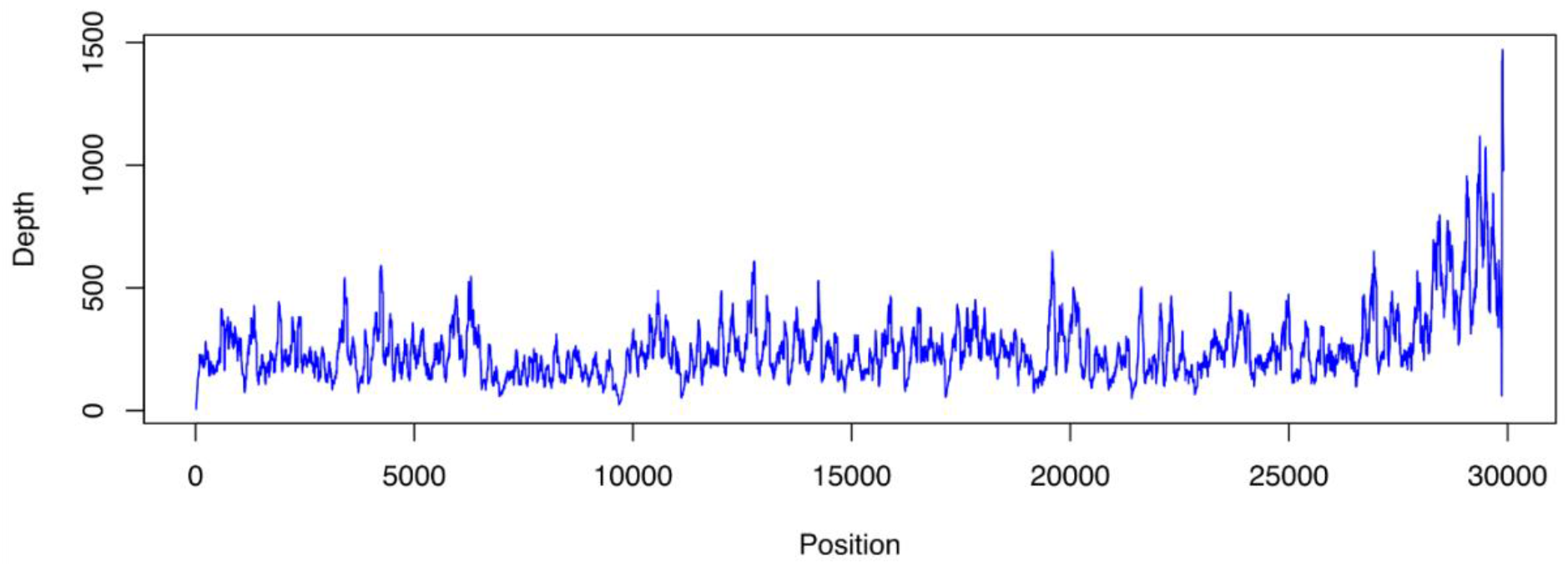
Depth of Coverage.

In addition, we also *de novo* assembled the reads with MEGAHIT software. Then, we performed a BLASTN analysis of the contigs using the NBCI non-redundant database. BLAST output showed that one contig of 29 861 base pairs hits the SARS-CoV-2 genome with a high percent identity. The hit corresponded to the SARS-CoV-2 genome with 100% coverage of the query and 99.96% of percent identity.

To compare the consensus sequence of the resequencing versus *de novo* assembled genome, we performed an alignment of the two sequences using BLASTN. The two sequences were identical with only one ambiguous nucleotide (Y: C or T) at the position 847 in the reference-based consensus sequence. Compared to the reference Wuhan Hu-1 genome, 19 SNPs were identified of which two were in the 5’ untranslated region (UTR) region, six were synonymous mutations, and 11 non-synonymous (amino acid substitutions). We also noted one mutation at the position 28833 (C to T) which falls in the “Charité_N_R” primer target (Table S1).

Twenty-one (21) Malian sequences provided by a German laboratory deposited on GISAID were retrieved and a multiple alignment was performed with MAFFT. Subsequently, phylogenetic tree was constructed with a neighbor joining method implemented in the *adegenet* package in R. Our newly sequenced genome clustered with the existing Malian samples sequenced abroad (Figure 3).

**Figure 3:**
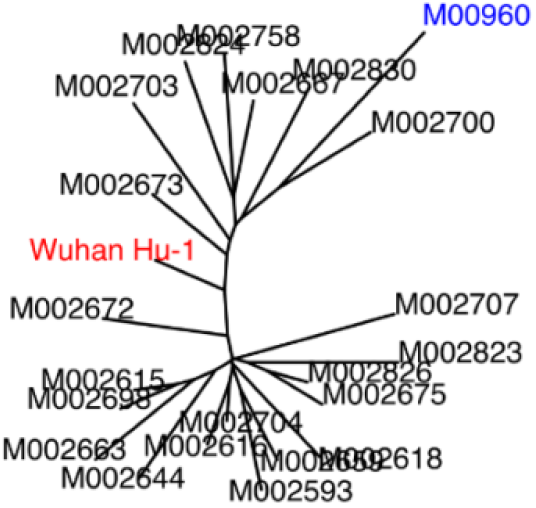
Neighbor Joining phylogenetic tree.

To determine the lineage of the new genome, we used the nextclade web application implemented on the Nextstrain website (https://nextstrain.org/sars-cov-2/) and the pangolin lineage assigner. The analysis assigns the genome to the B.1 lineage (pangolin classification), which is most common in USA, UK, France and it belongs to the 20A clade according to Nextstrain classification.

## Discussion

We report the first full-genome of SARS-CoV-2 sequenced and analyzed in Mali by Malian scientists. We have shown the feasibility of sequencing in resource limited settings. Genomic data are not only important for the development and surveillance of vaccines and diagnostic tests, but also essential for monitoring the evolution the virus and the transmission of the disease. Thus, our work has important implications for epidemiological tracing of the outbreak and could provide valuable information on routes of introduction of the virus and natural history of the SARS-CoV-2 circulating in Mali. Genomic data could help in identifying source and sink of the virus and guide interventions.

One of the major challenges during the pandemic is the availability of reagents. Therefore, we adapted an alternative solution using an agarose gel to estimate the size of our library. There was an overestimation of the size using the agarose gel. Even though the difference in library size estimation could be due to the adapters or clustering bias of shorter fragments, the library size estimation on agarose gel is less accurate. Nevertheless, the agarose gel electrophoresis could be useful for laboratories in resource limited setting with no access to expensive and sophisticated instruments for library size analysis such as a Bioanalyzer or a TapeStation or during a pandemic period when access to reagents becomes a challenge for some countries.

The recovered genome contained 19 SNP with some private mutations. Interestingly, the D614G mutation on the spike protein associated with increased infectivity and virion spike density [16]–[18]. The average number of cases in Timbuktu was the team in June 2020 was around 50 per day as compared to a national average case of 20 per day [19]. The presence of this mutation in Timbuktu might explain the sharp increase of cases in Timbuktu in June 2020 that has prompted the Malian Ministry of Health to send a team with a mobile laboratory to detect cases and trace contact to contain the outbreak.

The virus was likely been introduced in Mali from Europe as the lineage tracing showed that it belongs to the B1 lineage which is common in Europe and the USA. This is not surprising because of the frequent travels between Mali and Europe.

The sequencing platform was initially setup for malaria research. However, the COVID-19 pandemic has shown the versality of its use and underscores the importance to develop local sequencing capacity. The study demonstrated how an existing research infrastructure for a parasitic diseases can be leveraged to tackle a global health problem.

Our study has some limitations. One of these was the lack of quality control of total RNA before library preparation. For instance, we could not check the quality of the total RNA and did not know if the RNA was degraded or not because we did not have all the reagents required to do so. Decision to sequence the sample was solely based on the quantification of total RNA. Nonetheless, during this pandemic period, access to reagents remains a real challenge, quality control of the total RNA may be bypassed. Another issue was the library size estimation accuracy. Even though, an agarose gel analysis represents an alternative solution to overcome fragment size determination with expensive equipment it may overestimate the fragment size.

In summary, we provide the first genome of SARS-CoV-2 sequenced locally in Mali. Our result set a precedent for genomic surveillance of strains circulating in the country. We also offer the sequencing platform that can be used by scientists from neighboring countries. The platform could be used to provide insights in tracking the SAR-CoV-2 transmission dynamics, presence and evolution of variants and data generated could inform health authorities with local data for a better control of the disease.

## Data availability

Genome sequence has been deposited on the GISAID repository under the accession number EPI_ISL_683835 (https://www.gisaid.org/).

## Acknowledgments

- We are thankful to Africa CDC and Illumina for providing us with reagents.
- French Development Agency (AFD) supported the acquisition of the Illumina MiSeq instrument
- This work was supported through the DELTAS Africa Initiative [DELGEME grant 107740/Z/15/Z]. The DELTAS Africa Initiative is an independent funding scheme of the African Academy of Sciences (AAS)’s Alliance for Accelerating Excellence in Science in Africa (AESA) and supported by the New Partnership for Africa’s Development Planning and Coordinating Agency (NEPAD Agency) with funding from the Wellcome Trust [DELGEME grant 107740/Z/15/Z] and the UK government. The views expressed in this publication are those of the author(s) and not necessarily those of AAS, NEPAD Agency, Wellcome Trust or the UK government

## Conflict of interest

Authors declared no conflict of interest.

## Notes

### Competing Interest Statement

The authors have declared no competing interest.

## References

1 A. A. Pater et al., “Emergence and Evolution of a Prevalent New SARS-CoV-2 Variant in the United States,” bioRxiv, 2021.

2 M. Makoni, “South Africa responds to new SARS-CoV-2 variant,” Lancet (London, England). 2021.

3 J. W. Tang, P. A. Tambyah, and D. S. Hui, “Emergence of a new SARS-CoV-2 variant in the UK,” Journal of Infection. 2021.

4 A. Rambaut et al., “Preliminary genomic characterisation of an emergent SARS-CoV-2 lineage in the UK defined by a novel set of spike mutations,” Virological, 2020.

5 A. N. Happi, C. A. Ugwu, and C. T. Happi, “Tracking the emergence of new SARS-CoV-2 variants in South Africa,” Nature Medicine. 2021.

6 F. Naveca, V. Souza, and C. Costa, “COVID-19 epidemic in the Brazilian state of Amazonas was driven by long-term persistence of endemic SARS-CoV-2 lineages and the recent emergence of the new Variant of Concern P. 1,” Res. Sq. Sq., 2021.

7 P. C. Resende et al., “Spike E484K mutation in the first SARS-CoV-2 reinfection case confirmed in Brazil,” Oxford Journals, 2021.

8 A. Okwuraiwe, C. Onwuamah, O. Amoo, and B. Salako, “First African SARS-CoV-2 genome sequence from Nigerian COVID-19 case,” Res. gate, 2020.

9 R. Ziraldo, M. J. Shoura, A. Z. Fire, and S. D. Levene, “Deconvolution of nucleic-acid length distributions: a gel electrophoresis analysis tool and applications,” Nucleic Acids Res., vol. 47, no. 16, pp. e92–e92, Sep. 2019.

10 H. Li and R. Durbin, “Fast and accurate short read alignment with Burrows-Wheeler transform,” Bioinformatics, 2009.

11 H. Li et al., “The Sequence Alignment/Map format and SAMtools,” Bioinformatics, 2009.

12 H. Thorvaldsdóttir, J. T. Robinson, and J. P. Mesirov, “Integrative Genomics Viewer (IGV): High-performance genomics data visualization and exploration,” Brief. Bioinform., 2013.

13 D. R. Cavener, “Comparison of the consensus sequence flanking translational start sites in Drosophila and vertebrates,” Nucleic Acids Res., vol. 15, no. 4, pp. 1353–1361, 1987.

14 D. Li, C.-M. Liu, R. Luo, K. Sadakane, and T.-W. Lam, “MEGAHIT: an ultra-fast single-node solution for large and complex metagenomics assembly via succinct de Bruijn graph,” Bioinformatics, vol. 31, no. 10, pp. 1674–1676, May 2015.

15 G. Marçais, A. L. Delcher, A. M. Phillippy, R. Coston, S. L. Salzberg, and A. Zimin, “MUMmer4: A fast and versatile genome alignment system,” PLOS Comput. Biol., vol. 14, no. 1, p. e1005944, Jan. 2018.

16 L. Yurkovetskiy et al., “SARS-CoV-2 Spike protein variant D614G increases infectivity and retains sensitivity to antibodies that target the receptor binding domain,” bioRxiv, 2020.

17 L. Zhang et al., “SARS-CoV-2 spike-protein D614G mutation increases virion spike density and infectivity,” Nat. Commun., 2020.

18 L. Zhang et al., “The D614G mutation in the SARS-CoV-2 spike protein reduces S1 shedding and increases infectivity,” bioRxiv. 2020.

19 M. de la S. Mali, “Institut National de Sante Publique,” 2020. [Online]. Available: https://insp.ml/covid-ml/.

