## Supplemental file for "Sequencing SARS-CoV-2 in a malaria research laboratory in Mali, West Africa: the road to sequencing the first SARS-CoV-2 genome in Mali" (from another version of this preprint: version 1, 2021-05-05)

**Supplemental materials**

Preprocessing of raw data (FASTQC)

de novo assembly with MEGAHIT

Alignment of SARS-COV-2 reference genome MN908947.3--bwa

Post-aligment processing of bam file with samtools and vcfR (% of aligned reads, 99% genome coverage, read depth ~250, genome size = 29kb)

Bam file visualized with IGV and consenus sequence

Comparison of the outputs alignment consensus vers de novo assembled genome--nucmer

Figure S1: Bioinformatics analysis workflow.

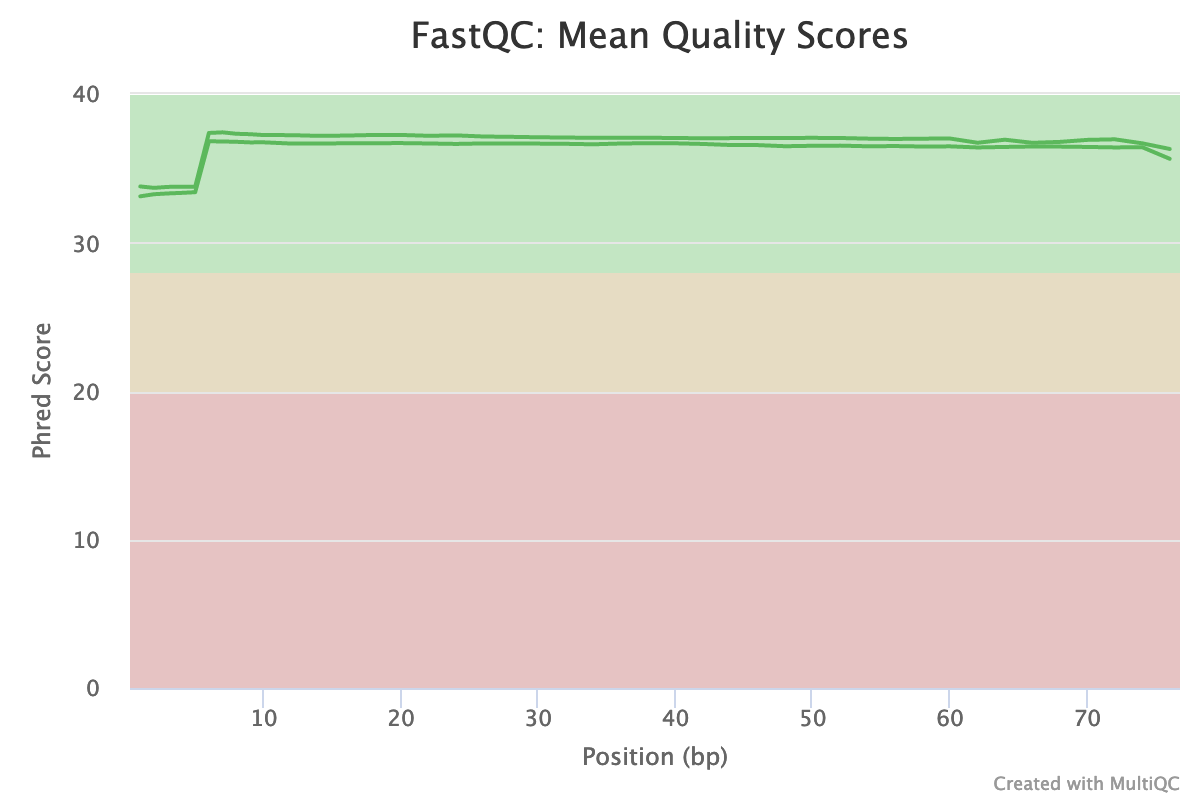

Figure S2: Read quality control

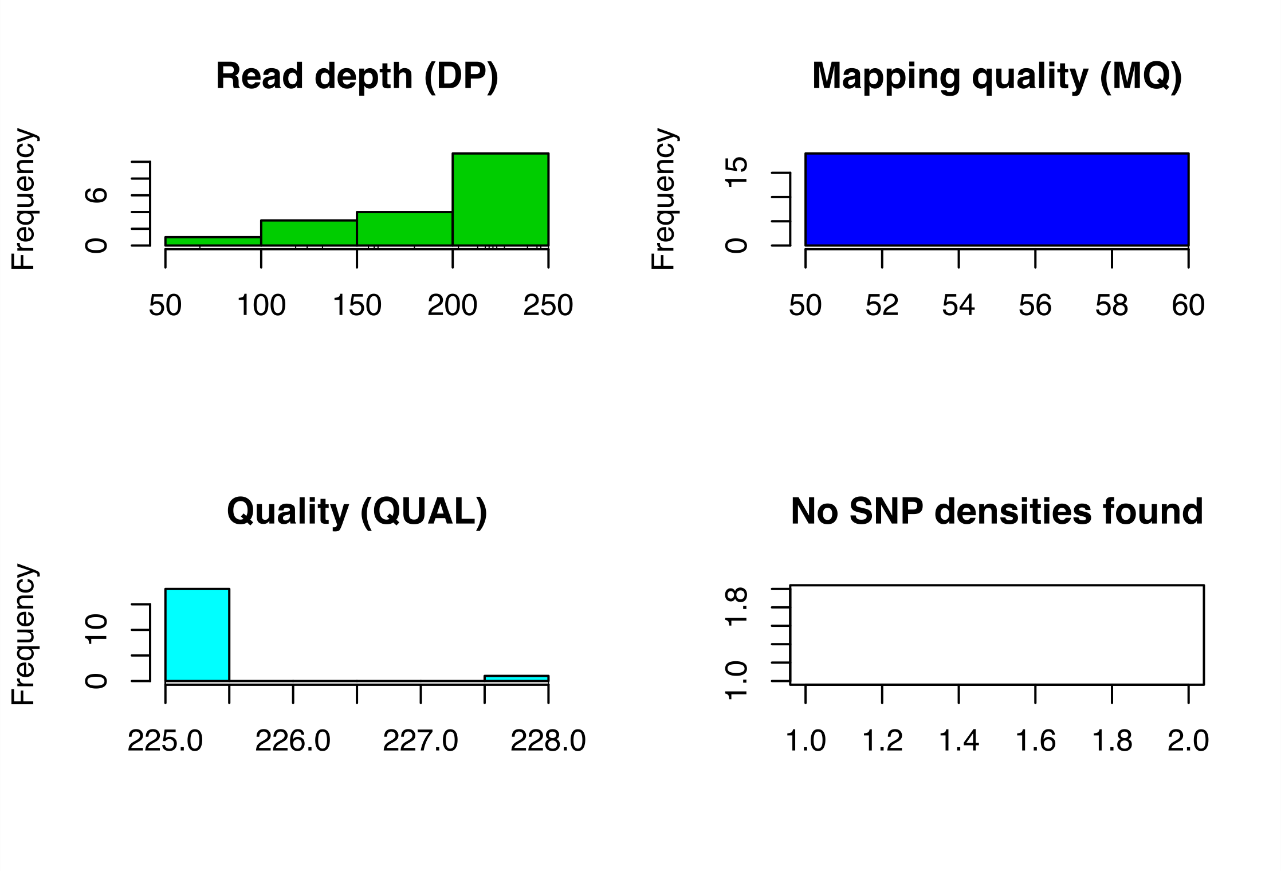

Figure S3: Alignment quality metrics

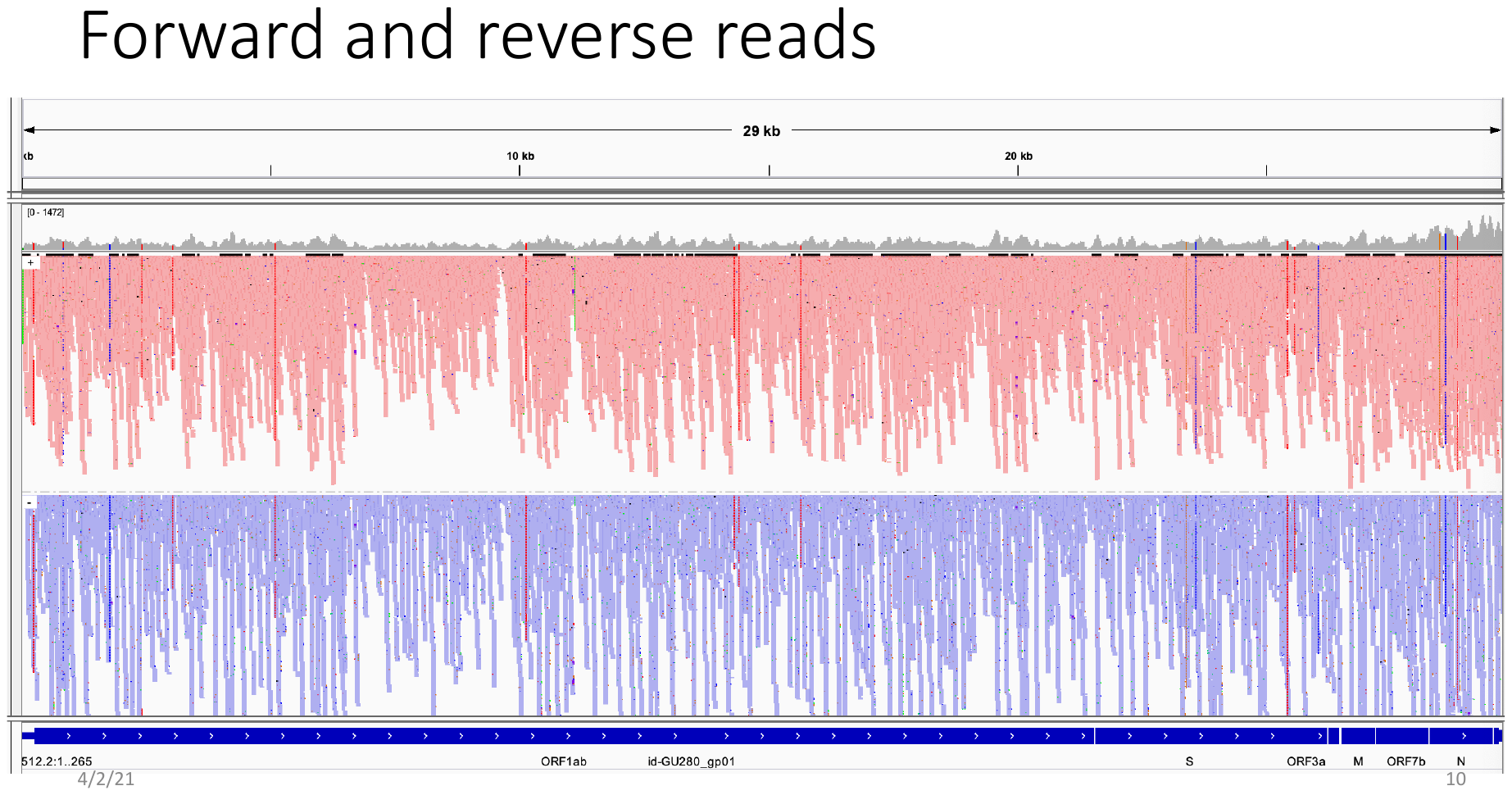

Figure S 4: Visualization of forward and reverse read alignment with IGV showing mutations supported by both reads.

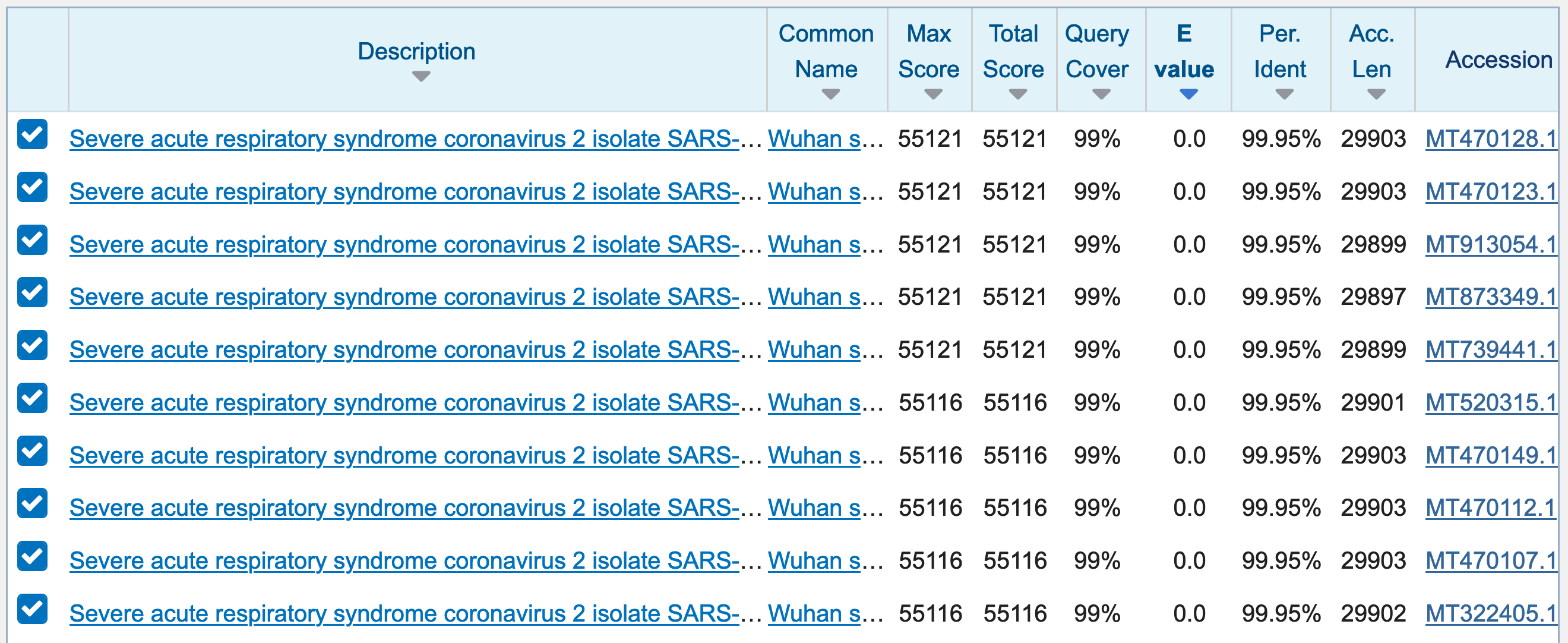

Figure S 5: Blast hits

Supplemental Table 1: SNPs in the new genome

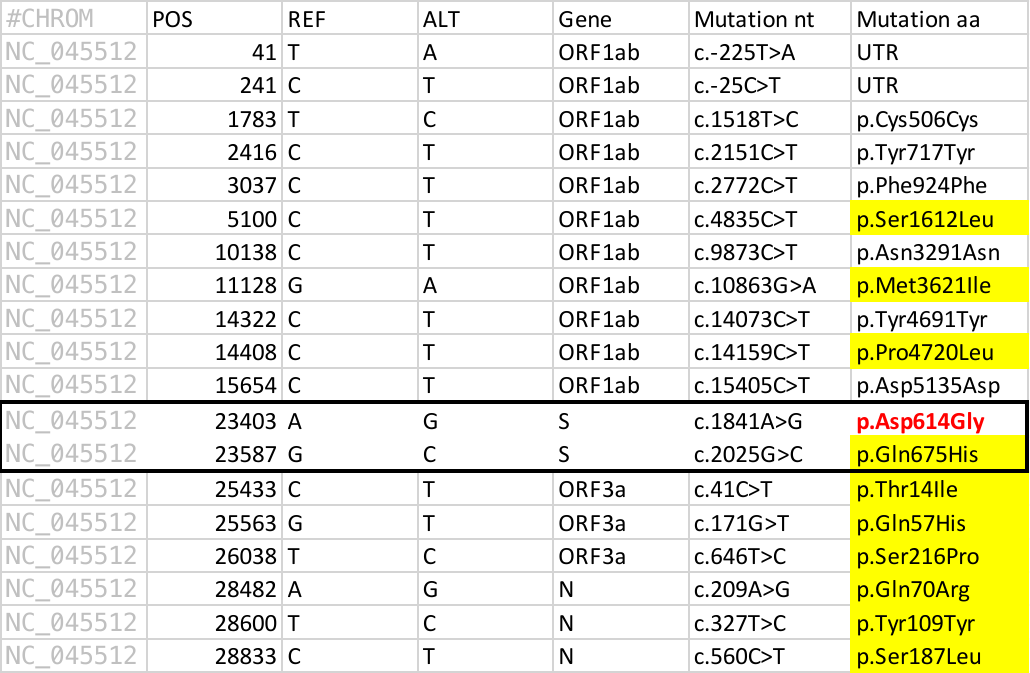
